# A unified framework links infant vulnerability with aging-related mortality dynamics

**DOI:** 10.64898/2026.05.05.722841

**Authors:** Ben Shenhar, Tzipi Strauss, Uri Alon

## Abstract

A central question in Geroscience is whether early-life mortality, which declines from birth to sexual maturity, and late-life mortality, which grows exponentially in time, can be understood within a shared conceptual framework. We show that stochastic threshold models can explain both phases by incorporating heterogeneity in neonatal vulnerability. Using U.S. National Center for Health Statistics data, we find that infant mortality risk is strongly associated with neonatal clinical markers such as Apgar scores, gestational age, and birth weight, suggesting that initial physiological differences persist across early life. We show that the ∼1/t mortality decline generically arises in stochastic threshold models via depletion of the most vulnerable, across a wide range of model specifications. Incorporating this mechanism into the Saturating-Removal model captures both the early decline and the later Gompertz acceleration, reproducing the full J-shaped mortality curve. Together, our findings link neonatal vulnerability to late-life mortality dynamics within a shared stochastic framework, supporting a life-course perspective on aging and longevity.

## Introduction

A key question in Geroscience is whether early-life human mortality is relevant to our understanding of the aging process. At first glance, early-life mortality behaves differently than late-life mortality - mortality declines approximately as 1/*t* over several decades of age from birth until sexual maturity, in contrast to the exponential rise of mortality that characterizes aging^1^. This J-shaped pattern has been well documented across organisms, including humans, primates, and other species^2,3^.

While the exponential increase in adult mortality has been extensively discussed^4–10^, early-life mortality has received much less attention, despite its commonality across the tree of life and over historical periods^2,3,11^. Establishing a mechanistic explanation for early-life mortality is therefore of interest, particularly if it can be understood within the same framework that accounts for the exponential rise in late-life mortality. Such a unified account would provide a coherent description of mortality from birth to death, and could offer insights into the determinants of aging.

Here, we provide a unified model of mortality across the life course. Using data from the U.S. National Center for Health Statistics (NCHS)^12,13^, we show that early-life mortality decline is consistent with vulnerability-dependent survival - mortality decreases because the most vulnerable die first, leaving a more robust population over time. We find that neonatal conditions, including the 5-minute Apgar score, birth weight, and gestational age, modulate mortality risk throughout the entire first year of life.

To model this, we extend the Saturating Removal (SR) model of aging dynamics to include vulnerable cohorts. The SR model is a biologically-calibrated equation for damage production and removal with stochastic noise^7^ (Methods). Its central assumption is that a dominant form of damage drives age-related changes. In the model, damage accumulates because production rate rises with age whereas removal saturates. Death occurs when damage crosses a death threshold. The model was originally inferred from longitudinal measurements of senescent cells in mice^7^ and has since been validated in additional species^14,15^ and against disease incidence statistics^16^. More recently, the SR model was used to reassess the heritability of human life expectancy^17^ and to explain the rigidity of the maximal human lifespan^18^.

To address early mortality, we introduce a vulnerable subpopulation with low death thresholds to the SR model. When these death thresholds are broadly distributed (uniform, power-law or mild exponential), the model recovers the characteristic 1/*t* decline of early-life mortality. We find that this behavior is robust to model details, arising in alternative stochastic threshold models and when recovery strength, rather than the death threshold, varies between individuals. We further show that the 5-minute Apgar score, a measure of neonatal condition, serves as a clinical correlate of the vulnerability parameter (death threshold). Among individuals dying at age t, both exhibit similar temporal dynamics, with mean values that depend approximately logarithmically on time.

Taken together, our results provide a ‘loss-of-the-most-vulnerable’ explanation for the 1/t decline in early-life mortality within a unified model that also captures late-life mortality and thus the entire J-shaped mortality curve. In this framework, vulnerability in early life influences survival across the lifespan, strengthening the case for a life-course perspective in Geroscience.

## Results

### Early-life mortality declines as 1/t from birth to sexual maturity

The human mortality curve is characteristically J-shaped, with a decline in childhood, a midlife plateau largely shaped by extrinsic hazards, and an exponential increase in adulthood (Fig. 1A). We explored the early-life mortality decline with contemporary mortality statistics from the United States^12,13,19^, United Kingdom (U.K.) ^20,21^, and Canada^22,23^, (Methods). Early-life mortality exhibits a log-log slope ranging from -1.01 to -1.12, (R^2>0.98, p<8e-6), consistent with previous studies^2,11,24,25^. We refer to this as 1/*t* behavior (Fig. 1B). This behavior spans 3.6 orders-of-magnitude, from 1 day of age to approximately 13 years. Using historical mortality data from the U.K., we find that despite improvements in life expectancy and living conditions^26^, this pattern has remained stable over the past century, with slopes ranging from -0.86 to -1.01 (R^2>0.98, p<1e-7) (Fig. 1C).

**Fig. 1.**
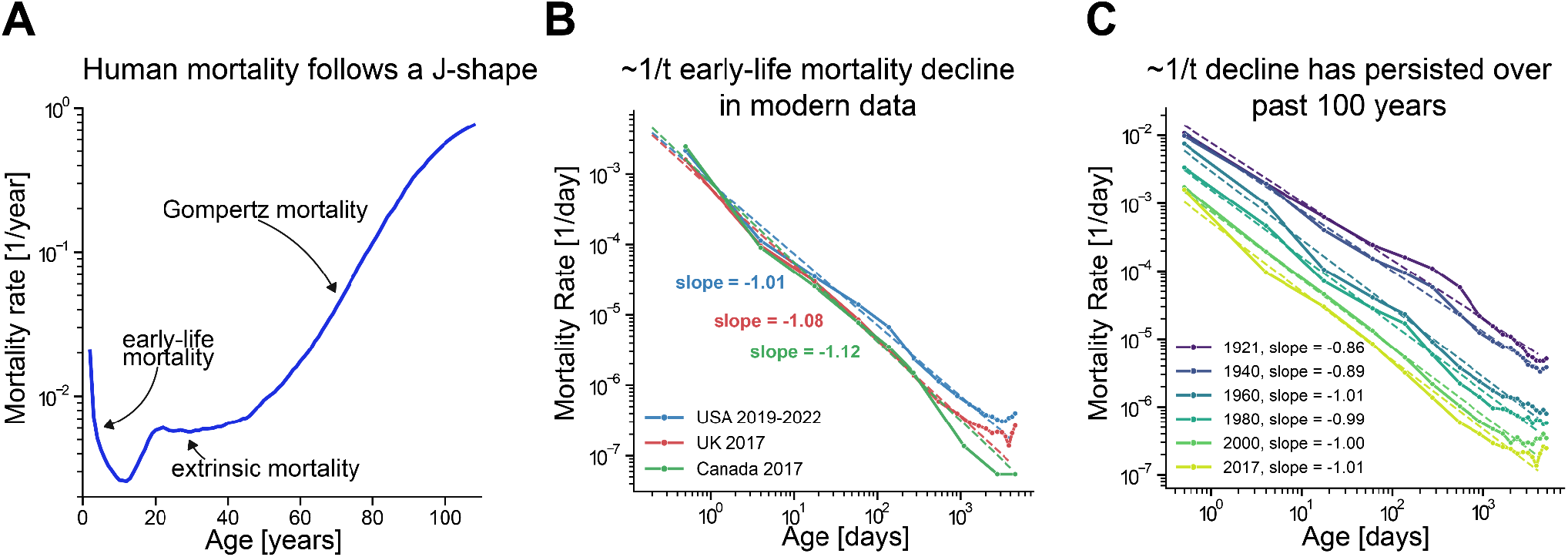
**(A)** Human mortality typically follows a J-shaped trajectory. Mortality declines through early life, then rises and plateaus due to extrinsic hazards, and finally increases exponentially in adulthood. Mortality rate shown for Sweden period data 1920, sexes pooled together. **(B)** Early-life mortality from birth to sexual maturity in datasets from the United States (2019-2022; blue), the United Kingdom (2017; red), and Canada (2017; green). All three show an approximately 1/t decline. **(C)** Historical early-life mortality in the United Kingdom from 1921 to 2017. Dashed lines indicate best-fit log–log slopes.

### Neonatal conditions are associated with elevated childhood mortality

To investigate the origins of early-life mortality, we analyzed the U.S. National Center for Health Statistics data^12^ on infant mortality during the first year of life to identify the major causes of death. Early-life mortality varies by cause, with roughly 50% of deaths attributable to perinatal conditions (Fig. 2A), primarily associated with preterm birth and low-birth weight (Fig. 2B). Approximately 20% of deaths are associated with congenital conditions, primarily cardiac and chromosomal abnormalities (Supp Fig. 1). Together, these findings suggest that impaired physiological function at birth is a major contributor to early-life mortality.

**Fig. 2.**
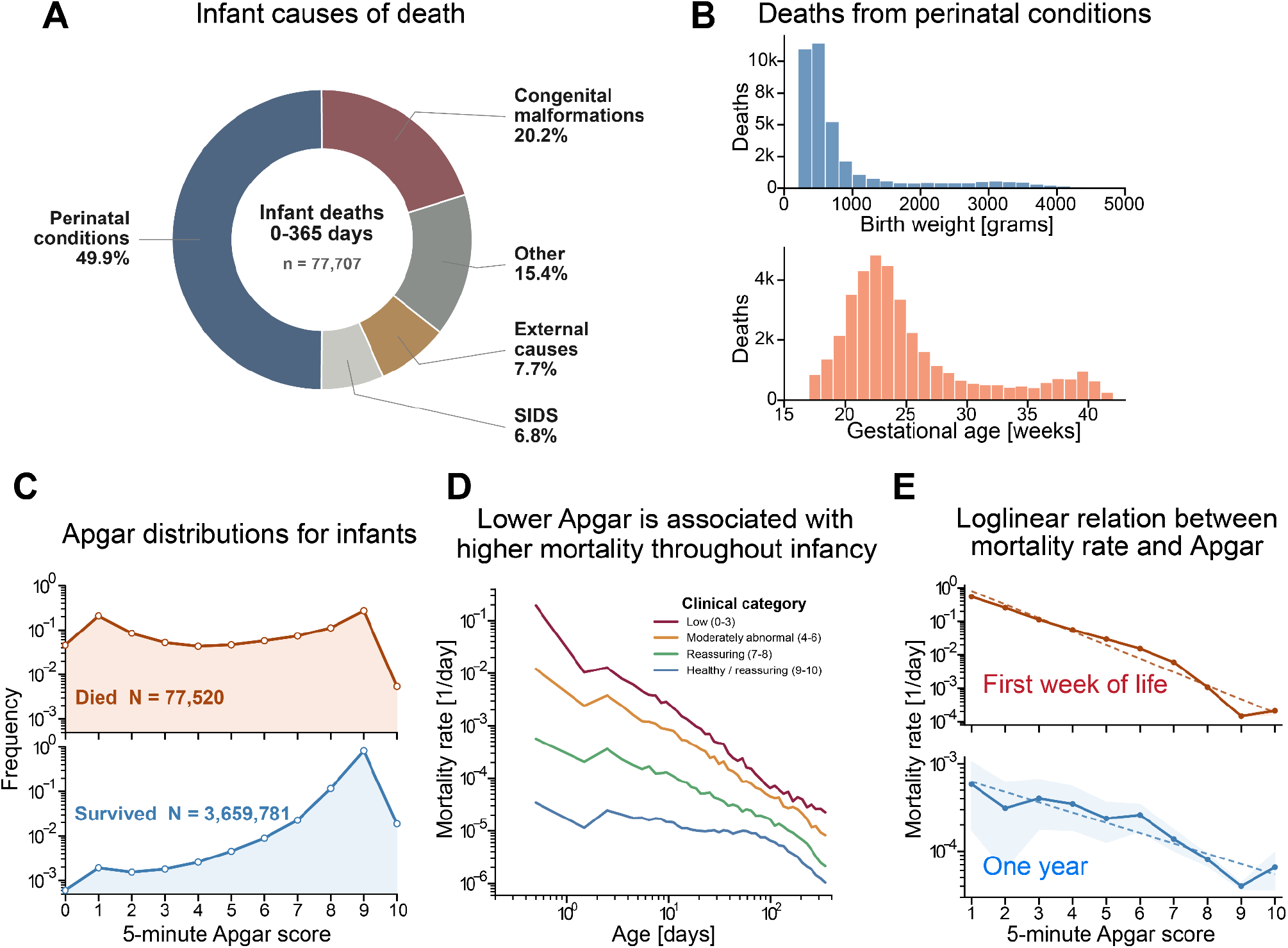
Early-life mortality risk is strongly dependent on perinatal conditions. **(A)** Distribution of causes of death in the first year of life in the U.S. National Center for Health Statistics (NCHS) linked birth/infant death files (2019-2022), grouped by ICD-10 category (Methods). SIDS refers to Sudden Infant Death Syndrome. **(B)** Birth weight (Upper) and gestational age (Lower) distributions for deaths due to perinatal conditions. **(C)** 5-minute Apgar score distributions (log-scale) for infants who died during the first year of life (Upper) and infants who survived (Lower). **(D)** Age-specific infant mortality rates stratified by 5-minute Apgar category at birth for U.S. infants born 2019-2022. Mortality is shown for clinical Apgar groupings 0–3, 4–6, 7–8, and 9–10. **(E)** Mortality rate as a function of 5-minute Apgar score evaluated at the first week of life (upper) and by the first birthday (lower). Shaded regions: bootstrap 95% CIs. First-week mortality (0–7 d); 1st birthday mortality (328.4–365 d).

We hypothesized that impaired function at birth is associated with persistently elevated mortality risk throughout early life. To test this, we analyzed data on 5-minute Apgar scores on millions of U.S. newborns from 2019-2022 (Methods). The Apgar score is a clinical assessment of neonatal condition at birth, based on five measures of physiological status each scored from 0-2: heart rate, respiration, reflex, muscle tone and appearance^27^. Summed scores range from 0 to 10, with higher scores indicating better conditions. Apgar distributions for infants who died during the first year of life were heavily centered around lower scores compared to surviving infants (Fig. 2C).

We stratified early-life mortality by Apgar score and found that lower scores were associated with higher mortality risk throughout the extent of the dataset, namely the entire first year of life (Fig. 2D). Risk of death during the first week of life is log-linear in Apgar score, such that log(m) = a_1_ Apgar + a_2_, with a_1_ = −0.93 ± 0.06, a_2_ = 0.72 (R^2^ = 0.96, p = 5 × 10−7). Remarkably, a similar trend was also found for risk of death at the 1st birthday, with log(m) = a_1_ Apgar + a_2_, with a_1_ = −0.27 ± 0.04, a_2_ = −7.1 (R^2^ = 0.87, p = 8 × 10−^5^) (Fig. 2E).

Impaired function at birth is thus linked to persistently elevated risk, even after the immediate postnatal period. Similar findings hold when stratifying by birth weight or gestational age (Supp. Fig. 2).

### The Saturating-Removal model with a subset of vulnerable individuals captures the J-shaped mortality curve

Because impaired neonatal conditions predict elevated mortality risk, we hypothesized that differential vulnerability drives the decline in mortality during early life. In this view, the most vulnerable tend to die first, leaving a progressively less vulnerable population over time with reduced mortality.

To test this idea, we asked whether the SR model, which offers a mechanistic explanation for the Gompertz phase of mortality^7^, can also account for the early-life regime. In the SR model, damage follows a stochastic trajectory, and death occurs upon first passage across a death threshold, Xc (Fig. 3A). We reasoned that preterm birth and congenital disorders may be modelled by reduced death thresholds (see next section). For this purpose, we interpret the SR model as a process in which “damage” x in infants represents deviations from physiological stability, and the death threshold is the largest deviation that can be physiologically compensated for.

**Fig. 3.**
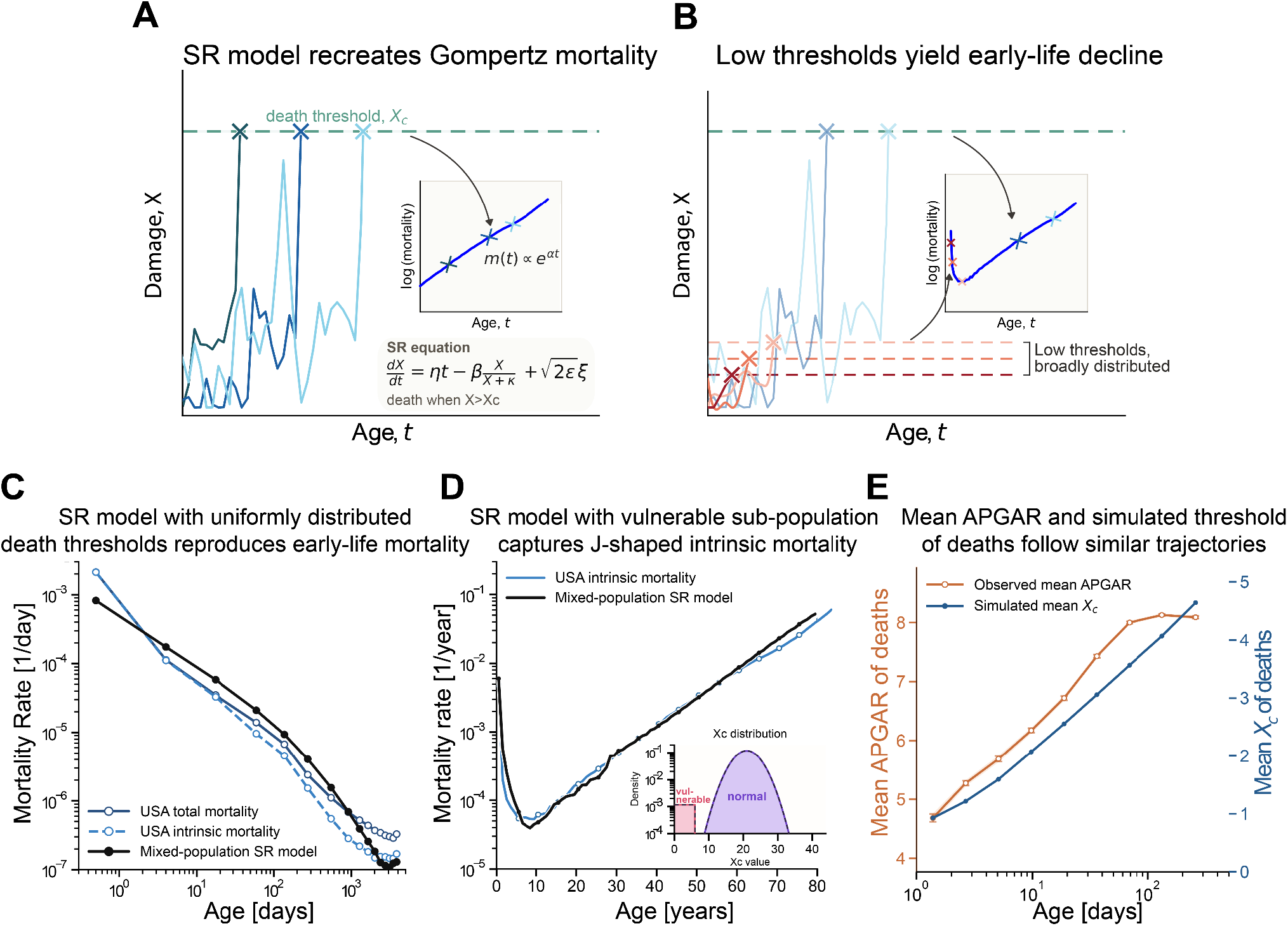
SR model with a heterogeneous vulnerable population recovers early-life mortality and J-shaped mortality. **(A)** Schematic of the SR model for a typical threshold population. Individuals accumulate damage X according to dX/dt = ηt − βX/(X + κ) + √(2ε)ξ and die when X reaches the robustness threshold Xc. The model yields an exponential increase in mortality in adulthood with slowdown at very old ages. **(B)** Schematic for a mixed population in which a small fraction of individuals have low-Xc thresholds, causing early deaths and an early-life mortality decline. **(C)** Early-life mortality comparison between U.S. data and the mixed-population SR model. **(D)** The same simulation compared against full-life, intrinsic mortality in the U.S. (Methods), exhibiting the full J-shaped mortality curve. Inset: The Xc threshold distribution used in the simulation. Red indicates the vulnerable subpopulation, purple indicates typical population. **(E)** Mean observed Apgar score and mean simulated Xc among those dying in matched age bins from the same simulation shown in D and E. Error bars throughout are SEM, often smaller than the marker size. The simulation used in all panels employed a threshold distribution centered at Xc = 21 with 15% Gaussian heterogeneity for the typical majority and a vulnerable subpopulation of weight w = 0.007 with *Xc* _*vulnerable*_ ∼ Uniform[0, 6]. The SR parameters are η = 4 × 10−^6^ d−^2^, β = 0.15 d−^1^, κ = 0.5, ε = 0.14 d−^1^.

We therefore consider a heterogenous population in the SR model, in which most individuals (>99%) have high death thresholds, and a small subpopulation has markedly lower thresholds. Individuals in the low-threshold group tend to die early, whereas those in the high-threshold group die in adulthood according to the Gompertz law (Fig. 3B).

We draw the vulnerable subpopulation death thresholds from a uniform distribution *Xc*_*vulnerable*_ ∼ *U*(0, *Xc*_*vul, max*_). We set the vulnerable-subpopulation weight to match the cumulative fraction of deaths before age 10 in U.S. 2021 period data, approximately 0.7%. The remainder of the population has high death thresholds drawn from a Gaussian distribution with parameters previously inferred from adult mortality data^15,18^.

We find both analytically and in simulations that a uniform low-threshold distribution leads to a 1/*t* decline of early-life mortality (Fig. 3C, see SI for analytical derivation). The simulation for the mixed-population vulnerable and non-vulnerable populations reproduces the J-shaped intrinsic mortality curve from birth to old age (Fig. 3D) with its declining and rising phases. Here, intrinsic mortality is defined via ICD-10 codes excluding external causes of death^28,29^ (Methods).

We next asked whether the decline in mortality indeed reflects progressive loss of the most vulnerable. The model predicts that the mean death threshold among individuals dying at age t rises slowly (logarithmically) with age (see SI). We compared this trend to the mean Apgar score of individuals dying at age t in the NCHS dataset (Fig. 3E). The data show a similar logarithmic increase, as quantified by a weighted linear regression of mean Apgar against log(age) (R^2^ =0.95, p = 5×10^-5^). This trend persists until saturation at approximately 100 days, reflecting the upper bound of the Apgar score (10), whereas the model’s death threshold continues to increase. These findings support the interpretation that early-life mortality decline arises from the progressive loss of the most vulnerable individuals.

We further examined the sensitivity of this behavior to the distribution of death thresholds within the vulnerable subpopulation. Power-law distributions reproduce nearly identical 1/t early-life mortality (SI, Supp Fig. 3). Very narrow distributions do not yield a 1/t behavior-for example, exponential distributions 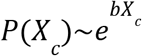 exhibit a mortality decline that goes as 1/*t*^1+*b*^ (SI), and are thus consistent with the data only with weak exponents (b ∼< 0.08). We conclude that the 1/t behavior is insensitive to the precise form of the threshold distribution and occurs generically for sufficiently broad distributions.

### Congenital malformations are consistent with reduced death threshold in adulthood

Our modelling approach treats congenital malformations as reduced death thresholds. We next asked whether this interpretation is consistent with survival patterns in individuals with congenital malformations who survive childhood. To test this, we analyzed the survival curve of patients with ventricular septal defects (VSD) that survived childhood using data from Eckerström et al.^16^ (Fig. 4A). The shape of the survival curve can indicate which SR model parameter differs from the control population, because each parameter change produces a distinct survival curve shape^18,30,31^. Reducing the death threshold Xc decreases mean lifespan more than maximal lifespan, yielding a shallow survival curve. In contrast, changes in production or removal rates affect both mean and maximal lifespan substantially, either proportionally or by a similar absolute shift, respectively.

**Fig. 4.**
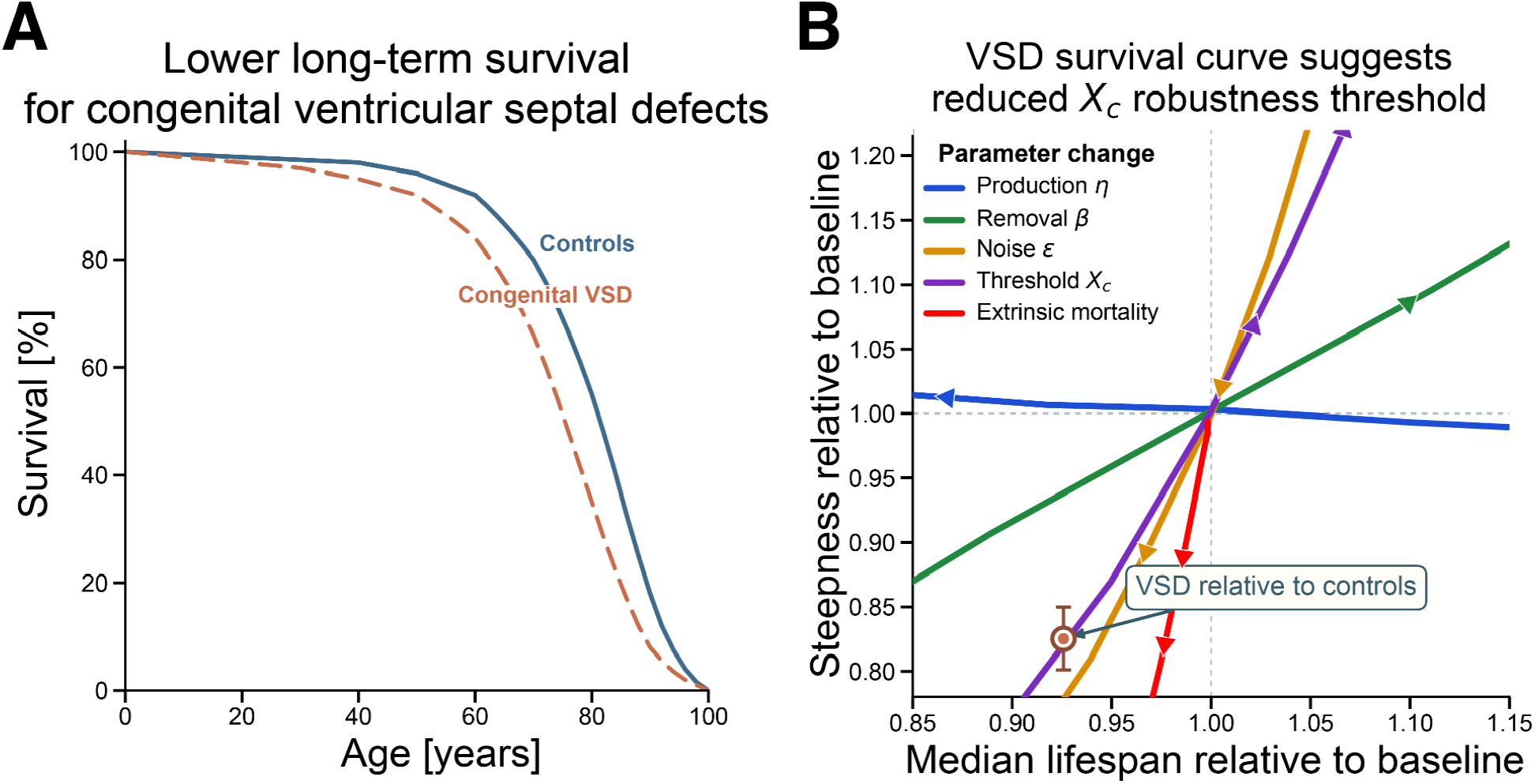
Congenital malformations are consistent with a low death threshold. **(A)** Survival curves for Danish patients with congenital ventricular septal defect (VSD; dashed; n = 9,136) and Danish controls from the same birth years (solid), reproduced from Eckerstrom et al.^32^ **(B)** Survival curve shape plane, relating changes in model parameters to changes in median lifespan and survival-curve steepness, defined as median/IQR. The annotated point is from the reproduced VSD survival curve relative to matched controls^32^. Error bars show bootstrap 95% confidence intervals computed from pseudo-lifespans sampled from the digitized survival curve.

These effects can be summarized in a steepness–longevity plot^18,30^, where changes in survival curve steepness (defined as median/IQR) are plotted against changes in median lifespan for different parameter shifts. The VSD survival curve (Fig. 4A) aligns with reduced threshold Xc (Fig. 4B). We therefore conclude that the VSD cohort is consistent with a reduced death threshold persisting into old age.

### 1/*t* early-life mortality decline is a generic property of stochastic threshold models with broad heterogeneity

We next asked whether the 1/t early-life mortality decline is specific to the SR model with heterogeneous death thresholds, or whether it also arises from other sources of heterogeneity and in other stochastic threshold models.

We first examined alternative sources of heterogeneity within the SR model. Assigning the vulnerable cohort a broad distribution of other model parameters also produced 1/t behavior, including heterogeneity in the damage-removal rate and in the noise amplitude (SI). Thus, the 1/t decline is robust to the specific biological origin of vulnerability, provided that vulnerability is broadly distributed.

We then considered a minimal stochastic threshold model distinct from the SR model, designed to capture only the early-life regime. In this model, x represents deviation from physiological stability. Biological regulation generates a restoring force that drives x back toward zero, while stochastic perturbations drive fluctuations away from zero. This is modeled as motion within a harmonic potential 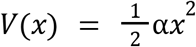, corresponding to the Ornstein-Uhlenbeck process^33^ (Methods, Fig. S4A). Death occurs when *x* crosses a critical threshold *X*_*c*_, corresponding to a deviation too large to be compensated for.

We considered two forms of vulnerability variation: (i) heterogeneity in the death threshold *X*_*c*_ and (ii) heterogeneity in the restoring force α (Fig. S4A). In both cases, the vulnerable individuals were assumed to constitute a small fraction of the total population (see SI). As in the SR model, a uniform distribution of restoring forces produced an approximately 1/*t* mortality. For a uniform distribution in barrier distances, the analytical result is 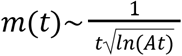, which for *At* ≫ 1 may be approximated as 1/*t* over the observed range (SI, Fig. S4B).

These results suggest that the 1/t early-life mortality decline is a generic feature of stochastic threshold models of physiological stability. Mechanistically, it arises from the progressive depletion of the most vulnerable individuals when vulnerability is broadly distributed. When implemented within the SR model framework, this mechanism recovers both the early-life mortality decline and the later Gompertz acceleration.

## Discussion

This study links early-life mortality decline with late-life mortality increase within the same mechanistic aging framework. The early-life decline arises from the progressive loss of the most vulnerable individuals, leaving a more robust surviving population with lower average mortality risk. Indicators of increased initial vulnerability, including low Apgar scores, prematurity, and low birth weight, all predict elevated mortality during the first year of life. In this context, early-life mortality should not be viewed solely as acute pathology, but as a manifestation of differences in developmental robustness at birth.

We showed that the 1/t decline is captured by the SR model by modeling a vulnerable subpopulation with broadly distributed death thresholds. When combined with the larger non-vulnerable cohort, the SR model offers a unified description of the full J-shaped mortality curve across the human lifespan. The early-life decline is insensitive to model details: heterogeneity in other vulnerability-related parameters, such as damage removal or noise amplitude, produces similar behavior. We further find that 1/t mortality dynamics also arise in a minimal stochastic threshold-crossing model, indicating that this behavior is a generic consequence of progressive loss of the most vulnerable when vulnerability is broadly distributed - uniform, power-law, or mildly exponential. In each case, early deaths selectively remove the most vulnerable individuals, increasing the mean robustness of the surviving population.

We further found that the neonatal Apgar score may serve as a clinical proxy for the effective death threshold of the vulnerable subpopulation. The mean Apgar score of individuals dying at age t increases logarithmically with age, consistent with the predicted increase in the mean death threshold among surviving individuals in the SR model.

A broad distribution of vulnerability may arise from several underlying biological mechanisms. Congenital abnormalities vary widely in size and severity^34,35^, which could generate a broad distribution of effective robustness thresholds. Ventricular septal defects provide one example, in which mild and moderate defect sizes occur at comparable frequencies^34,35^. Low Apgar score distributions for various neonatal conditions are also broad^36^. Nonetheless, this interpretation of broad vulnerability distribution remains suggestive, and the mapping from clinical heterogeneity to an idealized threshold distribution requires further investigation.

A limitation of the present study is that vulnerability was modelled as time-independent within individuals. Even after stratifying by initial condition, mortality continues to decline during childhood, which is consistent with robustness increasing with age. In reality, vulnerability is dynamic. It evolves rapidly during infancy as organ systems mature, physiological regulation stabilizes and adaptive capacity increases. Extending the model to include time-dependent thresholds or other forms of developmental change is therefore a direction for future work. In addition, early-life survival reflects not only intrinsic physiological vulnerability but also its interaction with the clinical environment, including the level of neonatal care and medical support, which are not explicitly captured in the current model.

In summary, early-life mortality can be understood within the same mechanistic aging framework as old-age mortality. A loss-of-the-most-vulnerable dynamic explains the 1/*t* decline of mortality at early ages, a pattern that also appears in primates. Our findings support a shift from viewing early-life mortality as a transient phenomenon to understanding it within a life-course framework, in which early differences in physiological robustness may define trajectories of survival extending far beyond the neonatal period. From a clinical perspective, this study motivates a search for biomarkers of neonatal vulnerability, and further research into the predictive capacity of such early biomarkers for later-life mortality. Interventions that increase robustness early in life may thus have long-lasting effects on survival. Early life should be considered within a life-course perspective on Geroscience, linking developmental biology with aging and longevity.

## Materials and Methods

### Materials

#### Late-life Mortality Data Source

Period mortality data for the United States, the United Kingdom, and Canada, spanning from 1 year of age onwards, were obtained from the Human Mortality Database^37^ and pooled across sexes.

#### Early-life Mortality Data Source

##### U.S

For the United States, we used the National Center for Health Statistics (NCHS) period linked birth/infant death public-use files for 2019-2022, accessed through the National Vital Statistics System (NVSS) linked data pages and the Vital Statistics Online portal. These linked files provide one record per infant death, together with variables from the linked birth record. In total, n = 3,737,301 records were used. From these files we used:

1. age at death
2. underlying ICD-10 cause of death (Underlying_Cause_Code)
3. 5-minute Apgar score (APGAR_Score_5min)
4. birth weight
5. gestational age

For the U.S. childhood extension mortality data, we used the 2021 NCHS mortality multiple cause-of-death public-use file.

##### U.K

For England and Wales (labeled “U.K.”), we used the Office for National Statistics (ONS) Child and infant mortality (by year of death) dataset for infant deaths and live births in the first year of life, together with the ONS Deaths registered by single year of age, U.K. dataset for the childhood extension.

For the historical plots, we combined the ONS infant bins with England-and-Wales Human Mortality Database period data^37^ to extend the curves through age 13 years.

##### Canada

For Canada, we used Statistics Canada table 13-10-0713-01 for infant deaths and infant mortality rates by infant age group, and Statistics Canada table 13-10-0710-01 for child age-group mortality and death counts used in the age-group reconstruction.

### Methods

### Mortality rate construction

Mortality is represented as an age-specific hazard. On an interval of length Δ*t*_*j*_ that contains *d*_*j*_ deaths, and has *N* _*j*_ individuals alive at the start of the interval, the mortality rate is defined as 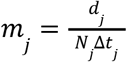. For plotting, interval-based points are placed at the midpoint of the interval. All intervals are left-inclusive and right-exclusive:

For U.K. and U.S. data, the intervals used were (in days): 0-1, 1-7, 7-28, 28-91, 91-183, 183-365, 365.25-730.50, 730.50-1095.75, 1095.75-1461.00, 1461.00-1826.25, 1826.25-2191.50, 2191.50-2556.75, 2556.75-2922.00, 2922.00-3287.25, 3287.25-3652.50, 3652.50-4017.75, 4017.75-4383, 4383-4748.25.

For Canada data, the intervals used were (in days): 0-1,1-7, 7-28, 28-91, 91-183, 183-365, 365.25-1826.25, 1826.25-3652.5, 3652.5-5478.75

### Extrinsic and intrinsic mortality rate construction

Intrinsic mortality rate constructions were derived for U.S. 2021 period data using ICD-10 underlying causes of death and Human Mortality Database (HMD) period death rates. Cause-of-death data were obtained from the U.S. 2021 Multiple Cause of Death microdata file.

For each age t, exposure-years were inferred as:

<preformate>

Exposure = all-cause deaths at age t / HMD mx

</preformate>

Deaths were partitioned into mutually exclusive categories according to ICD-10 underlying cause codes. Cause-specific mortality rates were then computed as:

<preformate>

cause-specific mx = deaths in partition / inferred exposure

</preformate>

By construction, this guarantees that intrinsic rate + extrinsic rate = HMD all-cause <monospace>mx</monospace> at every age and sex.

Extrinsic causes of death were defined using the following ICD-10 code groupings:

- V-Y: External causes and poisoning
- A-B: Infectious and parasitic diseases
- C33-C34: Trachea, bronchus, and lung cancer
- C53: Cervical cancer
- E40-E68: Nutritional deficiency and obesity disorders
- D50-D53, D59: Deficiency and acquired hemolytic anemias
- F10, F11-F19: Alcohol- and drug-related mental/behavioral disorders
- G00-G03: Meningitis
- I00-I09: Rheumatic fever and rheumatic heart disease
- J00-J99: Diseases of the respiratory system
- K70, K73-K74: Alcohol-related liver disease, chronic hepatitis, cirrhosis
- N10-N12, N30, N34-N39: Kidney and lower urinary tract infections/disorders
- N70-N77: Inflammatory diseases of the female pelvic/genitourinary tract
- O00-O99: Pregnancy, childbirth, and puerperium
- M35.3, M86: Polymyalgia rheumatica and osteomyelitis/infective bone disease Intrinsic mortality was defined by deaths from all other ICD-10 code groupings.

### SR Model

We used the saturating-removal (SR) mathematical model of aging dynamics, and expanded it here to simulate heterogeneous cohorts. The SR model describes damage X as a balance of production, removal, and noise:

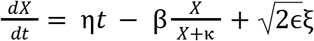

Damage production rate rises linearly with age (production = ηt) and damage is removed by Michaelis-Menten-like saturation kinetics, removal = βx/(κ+x), to represent removal mechanisms that become saturated at high damage levels. The parameters are β the maximal removal rate, κ the half-saturation constant, η the production parameter and ϵ the amplitude of Gaussian white noise ξ.

The full-life J-shaped mortality curves were generated by simulating a heterogeneous cohort with individual-specific death thresholds Xc. A small, vulnerable subpopulation (weight w=0.007) was assigned thresholds drawn from a uniform distribution on [0, *Xc*_*vul, max*_], representing early-life vulnerability. The remaining fraction (1-w) was assigned thresholds drawn from a Gaussian distribution, based on baseline parameters taken from prior work^15,18^. At initialization, each individual is assigned a fixed threshold and dies upon first-passage crossing of this threshold.

We simulate the model starting at *x*_0_ = 0 from t=0 to t=50,000 days with *n* = 10^6^, using an Euler-Maruyama scheme in Python with a reflecting boundary condition at x=0, dt = 0.02 days. We used a piecewise time-step (dt) scheme to resolve both early-life mortality, which varies on the scale of days, and late-life mortality, which on the scale of years: dt = 0.02 days for t ≤ 1, 0.05 days for 1 < t ≤ 10, 0.1 days for 10 < t ≤ 50, 0.5 days for 50 < t ≤ 100, 1.0 day for 100 < t ≤ 1000, 2.0 days for 1000 < t ≤ 5000, 4.0 days for 5000 < t ≤ 10000, and 8.0 days for 10000 < t ≤ 50000.

Mortality curves are calculated from the simulated lifespans of individuals using the mortality rate construction described above. The age intervals used match those employed in the analysis of U.S. and U.K. data.

### Minimal stochastic threshold model

The minimal stochastic threshold model that we consider is governed by Ornstein-Uhlenbeck dynamics. An over-damped particle undergoes diffusion in a harmonic potential 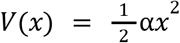 subject to Gaussian white-noise ξ, with diffusion coefficient ϵ:

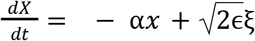

Death occurs as a first-passage time process when the trajectory reaches either of the symmetric absorbing boundaries ± *Xc*. Mortality rate is approximated as the first-passage-time distribution *f*(*t*), as early-life deaths are on the order of ∼10^-3^ of the total cohort: *S*_*cohort*_ (*t* _*early life*_) ≈ 1.

The stochastic dynamics are simulated using an Euler-Maruyama scheme, trajectories are initialized at *x*_0_ = 0, and simulated from t=0 to 10,000 days, with *n* = 5 × 10^4^. Simulations use a piecewise adaptive timestep, with starting dt < 10^−3^ /*α*_*max*_, and the largest dt < 5 10_−1_/*α*_*max*_ for times *t* > 1000 days. For the Xc barrier-distance distribution case, the parameter values are Xc ∼ U(1, 6.2), α = 1.6 day−^1^, and ε = 1 day−^1^. For the α restoring-force distribution case, the parameter values are α ∼ U(7.5, 300), Xc = 1.3 day−^1^, and ε = 3 day−^1^.

The first-passage-time distribution is computed from the simulated lifespans as the negative derivative of the Kaplan–Meier survival curve. It is evaluated on 72 logarithmically spaced bins between 0.1 and 10,000 days. Mean vulnerability parameters are then calculated within each bin.

### Cause-of-Death Grouping

U.S. infant causes of death were taken from Underlying_Cause_Code in the linked birth/infant death files. The top-level groups are:

- perinatal conditions: ICD-10 codes starting with P
- congenital malformations: ICD-10 codes starting with Q
- sudden infant death syndrome (SIDS): code R95
- external causes: ICD-10 codes starting with V, W, X, or Y
- other: all remaining codes

## Acknowledgements

Funding was provided by the European Research Council (grant agreement No. 856487) and by the Sagol Institute for Longevity Research, the Weizmann Institute of Science (B.S., U.A.). U.A. is the incumbent of the Abisch-Frenkel Professorial Chair.

## Supplemental Figures

**Supplemental Figure S1.**
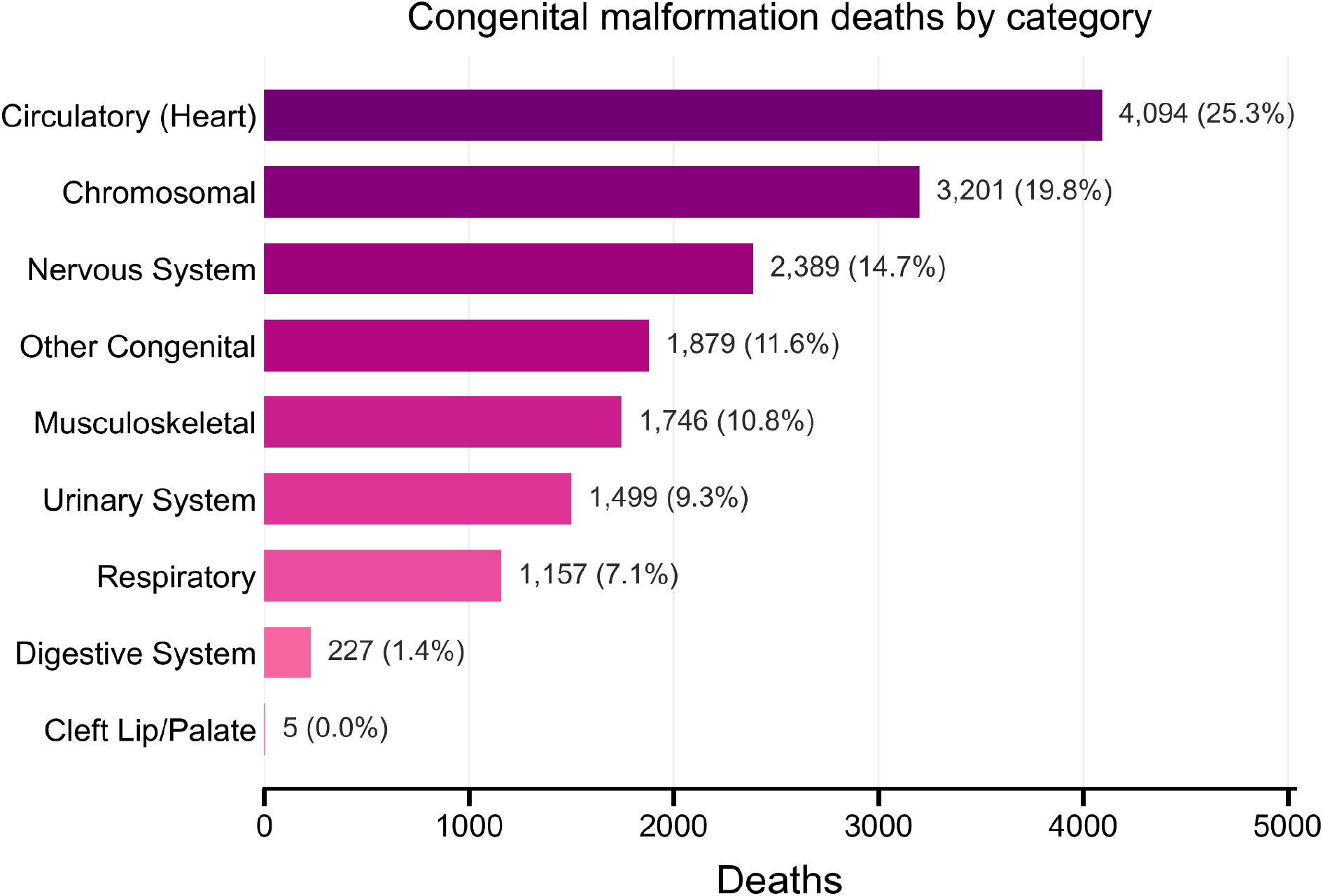
Distribution of congenital-malformation deaths by category in the United States. Infant deaths assigned to congenital malformations in the U.S. NCHS linked birth/infant death files, pooled across 2019-2022, grouped into broad ICD-10 congenital categories. Values indicate the number and fraction of deaths within the congenital-malformation subset.

**Supplemental Figure S2.**
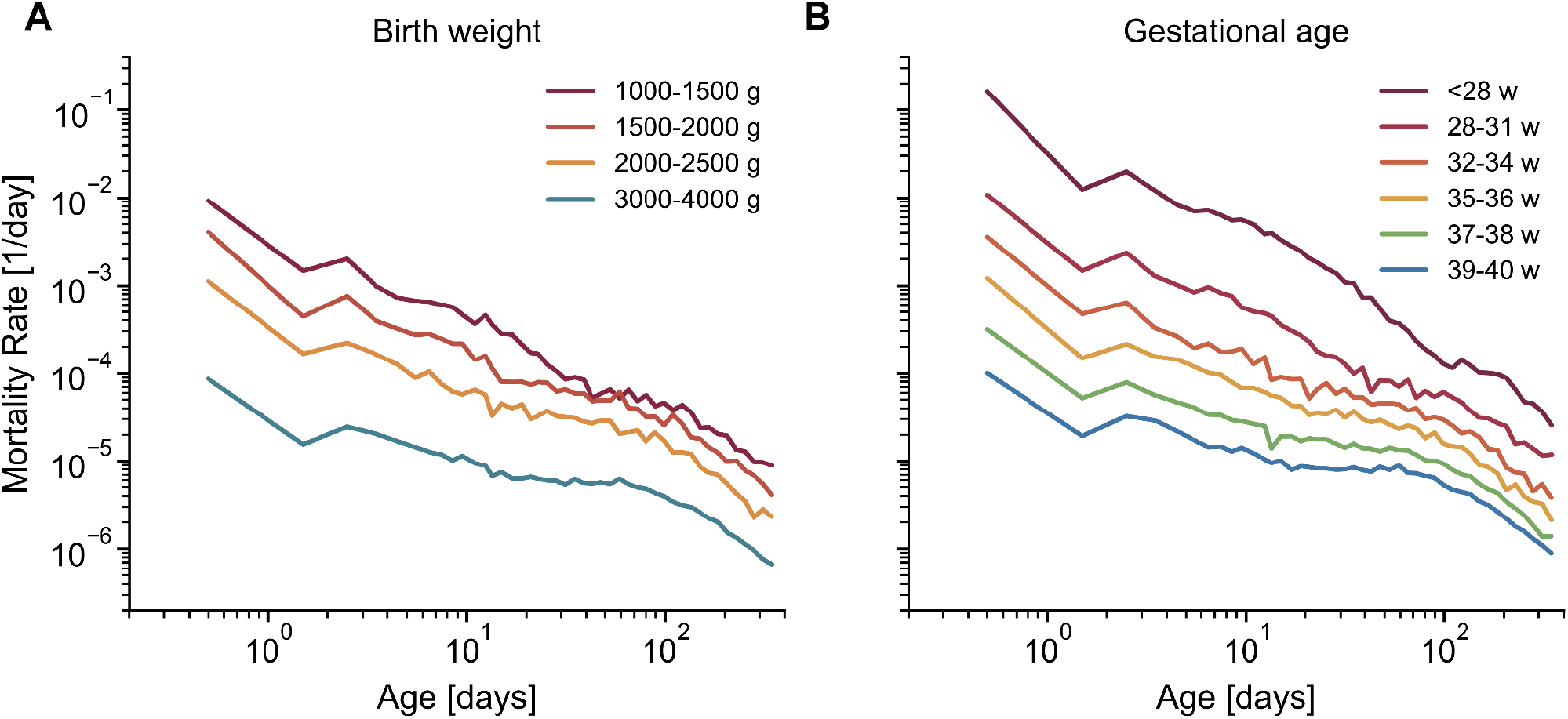
Infant mortality stratified by birth weight and gestational age in the United States. (A) Infant mortality in the U.S., stratified by birth weight. (B) Infant mortality in the U.S., stratified by gestational age. Curves were computed from pooled 2019-2022 NCHS linked birth/infant death data.

**Supplemental Figure S3.**
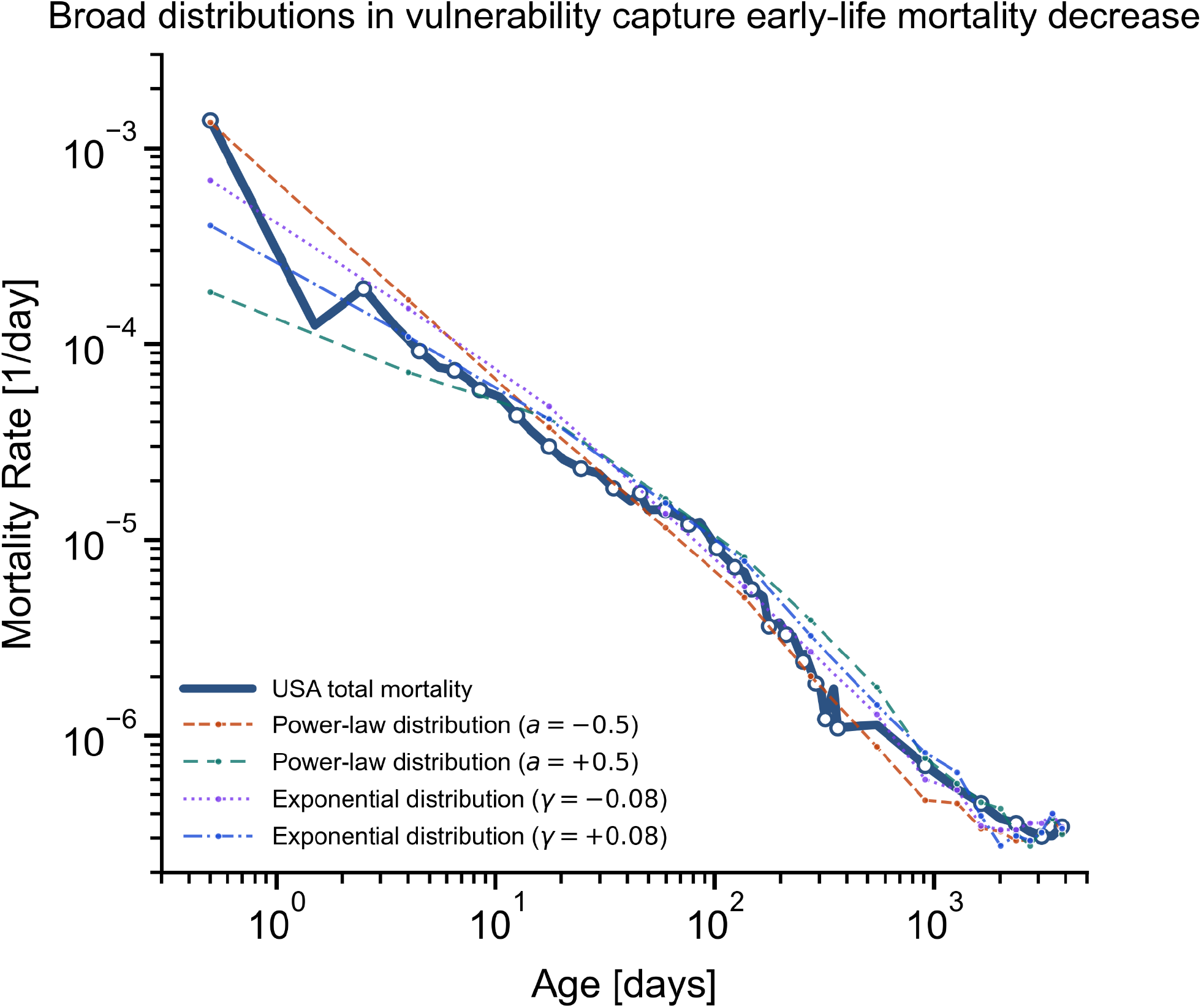
Broad distributions in death thresholds in the SR model reproduce early-life mortality decline. For a small vulnerable sub-population (weight w = 0.007), the death threshold Xc is drawn from truncated, normalized power-law distributions (exponent a = ±0.5) or exponential distributions (parameter γ = ±0.08) on the interval [0.05, 6], mixed with the normal Gaussian population. All other parameters match Figure 3. Curves show USA total mortality (data) and simulated mortality curves for each distribution.

**Supplemental Figure S4.**
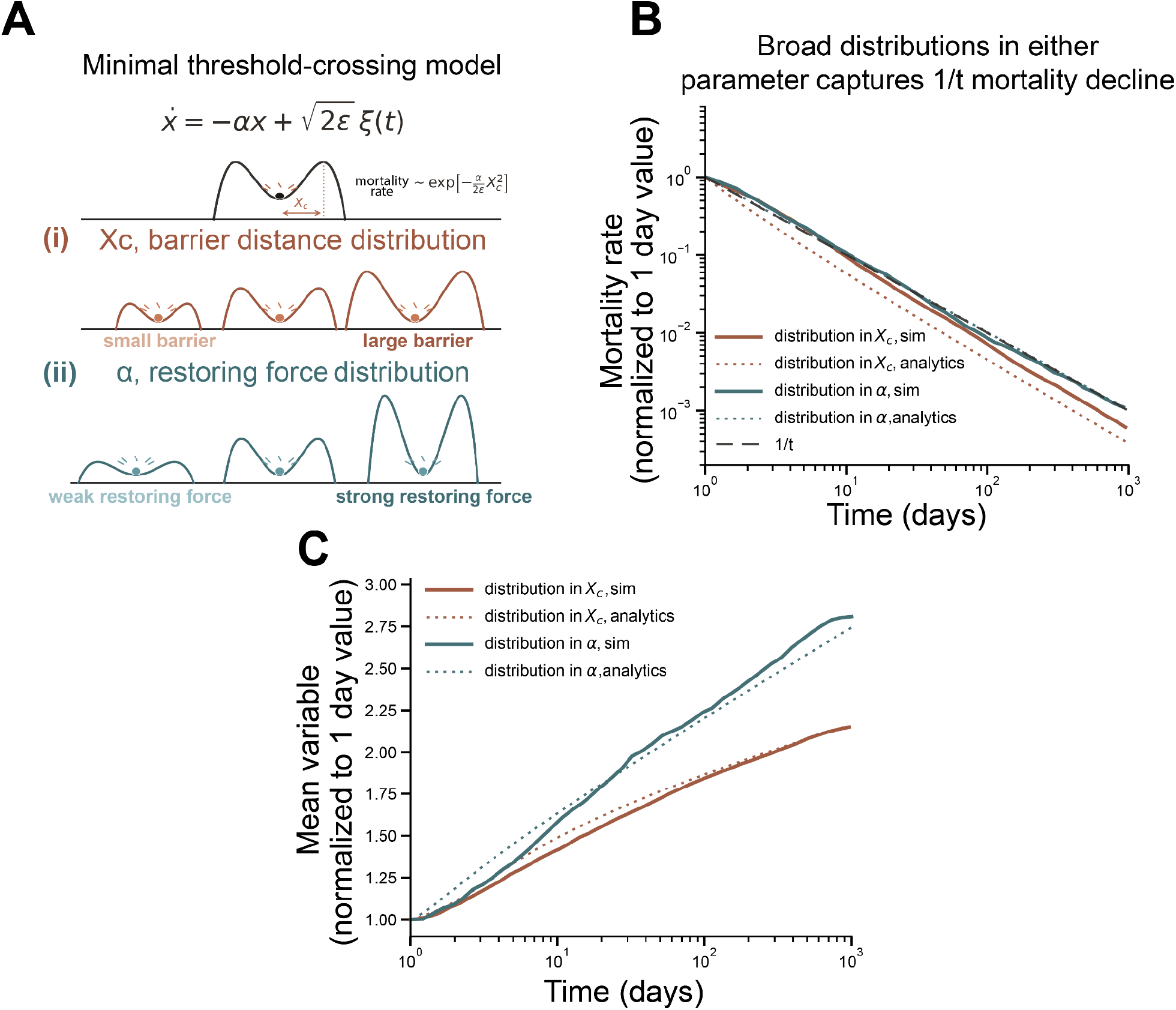
Heterogeneity in barrier distance or restoring force generates correct early-life mortality behavior in a general threshold model. **(A)** Schematic of the threshold model. Top: Stochastic trajectories of *x* in a harmonic well. α is the restoring force, ϵ sets the fluctuation amplitude, and ξ(*t*) is Gaussian white noise. Death occurs upon crossing the symmetric absorbing boundaries at ±Xc. Middle: Population heterogeneity is introduced through a uniform distribution in barrier distance Xc. Bottom row: Population heterogeneity is introduced through a uniform distribution in restoring force α. **(B)** Mortality rates, shown as the first-passage time density normalized to its value at 1 day, for populations with uniform distributions in barrier distance Xc (brown) or restoring force α (teal). Solid lines show simulation results; dotted lines show the corresponding analytic expectations. A dashed 1/t guide is shown for reference. **(C)** Mean vulnerability variable among individuals dying at each age, normalized to 1 day value. Solid lines show the mean Xc (brown) or mean α (teal) dying at time t; dotted lines show analytical predictions (see SI). Consistent with the theory, variation in restoring force produces an approximately *ln*(*t*) drift in the characteristic dying subpopulation, whereas variation in barrier distance produces a slower 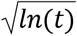 drift. Model parameters: barrier distribution case, *X* _*c*_ ∼ *U*(1. 0, 6. 2), α = 1. 6 [1/*day*], ϵ = 1 [1/*day*^2^]; restoring-force distribution case α ∼ *U*(7. 5, 300. 0), *X*_c_ = 1. 3, ϵ = 3 [1/*day*^2^ ].

