## Supplemental Information for "A unified framework links infant vulnerability with aging-related mortality dynamics"

**Abstract**

This Supplementary Information gives the analytical derivations used in the main text. We first derive general escape-rate and cohort-averaging results, then apply them to the SR model and to a generic Ornstein-Uhlenbeck model of physiological stability.

**S4 Summary of mortality scalings 10**

### S1 General escape-rate framework

#### S1.1 Escape-rate approximation for a steady-state stochastic process

Consider a particle fluctuating in a confining potential well,

$$\frac{\mathrm{d}X}{\mathrm{d}t} = -\nabla V(X) + \sqrt{2\epsilon}\,\xi,$$

where $\epsilon$ is the diffusion constant and $\xi$ is Gaussian white noise. Under the steady-state approximation, the probability density in the non-absorbing region is

$$P(X) = \mathcal{N} \exp\left[-\frac{V(X)}{\epsilon}\right],$$

where $\mathcal{N}$ is set by normalization over the bounded, non-absorbing region. With an absorbing boundary at $X = X_c$, the instantaneous escape rate is approximated by the probability flux at the boundary:

$$\lambda = \epsilon \frac{\mathrm{d}P(X)}{\mathrm{d}X}\bigg|_{X=X_c} = \mathcal{N}\epsilon\nabla V(X_c) \exp\left[-\frac{V(X_c)}{\epsilon}\right]. \tag{S1}$$

#### S1.2 Time-dependent cohort escape rate for a heterogeneous cohort

Let the instantaneous escape rate depend on a parameter $q$, with distribution $P(q)$ normalized as $\int P(q)\,\mathrm{d}q = 1$ over the allowed values of $q$. The parameter may represent, for example, a boundary location, a drift strength, or a potential-well parameter. For individuals with parameter value $q$, survival is

$$S_q(t) = \exp[-\lambda(q)t].$$

The cohort mortality rate is the survivor-weighted average of the parameter-specific escape rates:

$$m_{\mathrm{cohort}}(t) = \frac{\int \lambda(q)P(q)S_q(t)\,\mathrm{d}q}{\int P(q)S_q(t)\,\mathrm{d}q} = \frac{\int \lambda(q)P(q)\mathrm{e}^{-\lambda(q)t}\,\mathrm{d}q}{\int P(q)\mathrm{e}^{-\lambda(q)t}\,\mathrm{d}q}. \tag{S2}$$

The numerator is the cohort first-passage-time density,

$$f_{\mathrm{cohort}}(t) = \int \lambda(q)P(q)\mathrm{e}^{-\lambda(q)t}\,\mathrm{d}q,$$

and the denominator is the cohort survival function,

$$S_{\mathrm{cohort}}(t) = \int P(q)\mathrm{e}^{-\lambda(q)t}\,\mathrm{d}q.$$

##### S1.3 Small early-life mortality approximation

In the early-life regime considered here, only a small fraction of the cohort has died. Thus $S_{\text{cohort}}(t) \approx 1$, and (S2) is well approximated by the first-passage-time density:

$$m_{\text{cohort}}(t) \approx f_{\text{cohort}}(t). \tag{S3}$$

We use this approximation in the analytical derivations below.

##### S1.4 Evaporative-cooling analogy

The cohort expression has a useful analogy to statistical physics. If the escape rate is read as an energy, $\lambda(q) \to E(q)$, and time is read as inverse temperature, $t \to 1/(k_B T)$, then the cohort survival function has the form of a partition function,

$$S_{\text{cohort}}(t) \to \int P(q) \exp\left[-\frac{E(q)}{k_B T}\right] \mathrm{d}q.$$

The cohort mortality rate corresponds to the mean energy of the surviving population. At $t = 0$, the effective temperature is infinite and the mortality rate is dominated by the fastest escape rates. As time increases, the effective temperature decreases: high-escape-rate individuals are removed first, and the remaining cohort is progressively enriched for lower escape rates.

#### S2 SR model results

##### S2.1 Early-life approximation

The Langevin equation for the SR model is

$$\frac{\mathrm{d}X}{\mathrm{d}t} = \eta t - \beta \frac{X}{X + \kappa} + \sqrt{2\epsilon}\, \xi, \tag{S4}$$

where $\epsilon$ is the diffusion constant and $\xi$ is Gaussian white noise. The model has a reflecting boundary at $X = 0$ and an absorbing boundary at $X = X_c$. The characteristic timescale is $\tau = \frac{\beta}{\eta}$, the time at which stability is lost and $X$ begins drifting toward the absorbing boundary. Previous fits to human mortality data found $\tau \approx 100$ years. We focus on early-life mortality, $t \lesssim 13$ years. In this regime, $\eta t < \beta$, so the production term is negligible. Linearizing the removal term gives

$$\frac{\mathrm{d}X}{\mathrm{d}t} = -\beta + \sqrt{2\epsilon}\, \xi. \tag{S5}$$

##### S2.2 Instantaneous escape rate

For a Brownian particle with constant drift $-\beta$ starting at a reflecting boundary, the exact mean first-passage time (MFPT) to an absorbing boundary at $X_c$ can be obtained from the backward

Fokker-Planck equation:

$$\tau(X_c) = \frac{\epsilon}{\beta^2}\left[\exp\left(\frac{\beta X_c}{\epsilon}\right) - 1 - \frac{\beta X_c}{\epsilon}\right]. \tag{S6}$$

In the rare-event limit, $\beta X_c \gg \epsilon$, the exponential term dominates. The escape or mortality rate $\lambda = 1/\tau$ is therefore

$$\lambda(X_c) \approx \frac{\beta^2}{\epsilon}\exp\left[-\frac{\beta X_c}{\epsilon}\right]. \tag{S7}$$

#### S2.3 Uniform threshold distribution and $1/t$ mortality scaling

Assuming a uniform distribution of barrier heights from zero to a maximum threshold $X_{vul,max}$:

$$P_{\text{vulnerable}}(X_c) = \frac{1}{X_{vul,max}}, \qquad X_c \in [0, X_{vul,max}]. \tag{S8}$$

Below we show how this distribution produces a $1/t$ decline in early-life mortality.

##### S2.3.1 Frontier mapping

The frontier mapping gives an intuitive derivation. At observation time $t$, the deaths are dominated by the subpopulation whose characteristic lifetime satisfies $\tau \approx 1/\lambda \approx t$. The threshold currently being depleted satisfies

$$\lambda(X_c) = \frac{\beta^2}{\epsilon}\exp\left[-\frac{\beta X_c}{\epsilon}\right] \approx \frac{1}{t}.$$

Solving for the frontier threshold gives

$$X_c(t) = \frac{\epsilon}{\beta}\ln\left(\frac{\beta^2}{\epsilon}t\right).$$

Thus the mean barrier height among those dying moves logarithmically deeper into the threshold distribution. The frontier speed is

$$\frac{\mathrm{d}X_c}{\mathrm{d}t} = \frac{\mathrm{d}}{\mathrm{d}t}\left[\frac{\epsilon}{\beta}\ln\left(\frac{\beta^2}{\epsilon}t\right)\right] = \frac{\epsilon}{\beta t}.$$

The mortality rate is the density of thresholds at the frontier times the speed at which the frontier moves:

$$m(t) = P(X_c)\left|\frac{\mathrm{d}X_c}{\mathrm{d}t}\right| = \frac{\epsilon}{\beta X_{\max}}\frac{1}{t}. \tag{S9}$$

Because the escape rate in (S7) depends exponentially on $\beta X_c/\epsilon$, comparable heterogeneity in $\beta$ or $\epsilon$ can produce similar scaling behavior.

##### S2.3.2 Flux integral

The same result follows by explicitly integrating the death flux over the uniform distribution. Using (S3),

$$m_{\text{cohort}}(t) \approx \int_0^{X_{\max}} \frac{1}{X_{\max}} \left( \frac{\beta^2}{\epsilon} \exp\left[-\frac{\beta X_c}{\epsilon}\right]\right) \exp\left[-\frac{\beta^2}{\epsilon} t \exp\left(-\frac{\beta X_c}{\epsilon}\right)\right] \mathrm{d}X_c. \tag{S10}$$

Let

$$u = \frac{\beta^2}{\epsilon} t \exp\left[-\frac{\beta X_c}{\epsilon}\right].$$

Then

$$\begin{aligned} m_{\text{cohort}}(t) &= \frac{\epsilon}{\beta X_{\max} t} \int_{u_{\text{slow}}}^{u_{\text{fast}}} \mathrm{e}^{-u}\, \mathrm{d}u \\ &= \frac{\epsilon}{\beta X_{\max} t} \left(\mathrm{e}^{-u_{\text{slow}}} - \mathrm{e}^{-u_{\text{fast}}}\right), \end{aligned}$$

where

$$u_{\text{fast}} \equiv \frac{\beta^2}{\epsilon} t, \qquad u_{\text{slow}} \equiv \frac{\beta^2}{\epsilon} t \exp\left[-\frac{\beta X_{\max}}{\epsilon}\right].$$

Equivalently,

$$\begin{aligned} m_{\text{cohort}}(t) = \frac{\epsilon}{\beta X_{\max} t} \Bigg[ &\exp\left(-\frac{\beta^2}{\epsilon} t \exp\left[-\frac{\beta X_{\max}}{\epsilon}\right]\right) \\ &- \exp\left(-\frac{\beta^2}{\epsilon} t\right)\Bigg]. \end{aligned}$$

For human-fit parameter values, $\beta \approx \epsilon \approx 0.15\ \text{day}^{-1}$, so $t \gg \epsilon/\beta^2 \approx 6$ days implies $u_{\text{fast}} \to \infty$ and $\mathrm{e}^{-u_{\text{fast}}} \approx 0$. For the most stable frail modes near $X_c \sim X_{\max}$, the condition

$$t \ll \frac{\epsilon}{\beta^2} \exp\left(\frac{\beta X_{\max}}{\epsilon}\right)$$

implies $u_{\text{slow}} \to 0$ and $\mathrm{e}^{-u_{\text{slow}}} \approx 1$. In this intermediate-time regime,

$$m_{\text{cohort}}(t) \approx \frac{\epsilon}{\beta X_{\max}} \frac{1}{t}, \tag{S11}$$

matching (S9).

The power-law regime breaks down when

$$t \sim \frac{\epsilon}{\beta^2} \exp\left(\frac{\beta X_{\max}}{\epsilon}\right).$$

This condition constrains $X_{\max}$ because the empirical power-law decline extends through early childhood. For the mortality-data fit $X_{\max} = 6$, the breakdown time is approximately $6.7\mathrm{e}^6 \approx 2700$ days, or about 7.4 years. This is close to the observed end of the infant/childhood decline and well below the inferred healthy threshold $X_c \sim 21$ in the main-text fit.

#### S2.4 Additional threshold distributions

For non-uniform threshold distributions, the same frontier argument gives the leading mortality scaling. Define

$$A = \frac{\beta^2}{\epsilon}, \qquad X_c(t) = \frac{\epsilon}{\beta}\ln(At), \qquad \left|\frac{\mathrm{d}X_c}{\mathrm{d}t}\right| = \frac{\epsilon}{\beta t}.$$

Then

$$m(t) \approx P(X_c(t))\left|\frac{\mathrm{d}X_c}{\mathrm{d}t}\right|.$$

##### S2.4.1 Exponential threshold distribution

For

$$P(X_c) \propto \mathrm{e}^{\gamma X_c},$$

with a finite cutoff when $\gamma > 0$,

$$P(X_c(t)) \propto \exp\left[\gamma\frac{\epsilon}{\beta}\ln(At)\right] \propto t^{\gamma\epsilon/\beta}.$$

Therefore,

$$m(t) \propto t^{-1+\gamma\epsilon/\beta}. \tag{S12}$$

When $\gamma > 0$, mortality declines more slowly than $1/t$. When $\gamma < 0$, mortality declines faster than $1/t$.

##### S2.4.2 Power-law threshold distribution

For

$$P(X_c) \propto X_c^k,$$

with a finite cutoff when $k > 0$,

$$P(X_c(t)) \propto \left[\frac{\epsilon}{\beta}\ln\left(\frac{\beta^2}{\epsilon}t\right)\right]^k.$$

Thus

$$m(t) \propto \frac{\left[\ln\left(\frac{\beta^2}{\epsilon}t\right)\right]^k}{t}, \tag{S13}$$

up to a constant prefactor.

##### S2.5 Full mortality curve

To represent both the early-life decline and late-life aging, the full model uses a mixture of vulnerable and robust threshold distributions:

$$P_{\text{total}}(X_c) = wP_{\text{vulnerable}}(X_c) + (1-w)P_{\text{robust}}(X_c). \tag{S14}$$

The vulnerable component controls the early-life decline; the robust component controls the later Gompertz-like rise.

#### S3 Minimal stochastic threshold model: Ornstein-Uhlenbeck process

The Ornstein-Uhlenbeck process is

$$\mathrm{d}x = -\alpha x \ \mathrm{d}t + \sqrt{2\epsilon}\,\xi, \tag{S15}$$

with absorbing thresholds at $\pm X_c$.

##### S3.1 Instantaneous escape rate

Using the steady-state approximation from Section S1.1 and assuming a rare-event regime, we normalize the Boltzmann distribution over the real line. The escape rate is the total probability flux across the two absorbing boundaries,

$$J = \epsilon \left| \frac{\partial P}{\partial x} \right|_{x=X_c} + \epsilon \left| \frac{\partial P}{\partial x} \right|_{x=-X_c}.$$

This gives

$$\lambda = X_c \sqrt{\frac{8\alpha^3}{\pi\epsilon}} \exp\left(-\frac{\alpha X_c^2}{2\epsilon}\right). \tag{S16}$$

##### S3.2 Uniform distribution of restoring force $\alpha$

Assume a uniform distribution of restoring forces:

$$P(\alpha) = \frac{1}{\alpha_{\max}}, \qquad \alpha \in [0, \alpha_{\max}].$$

Writing the escape rate as

$$\lambda(\alpha) = C_1 \alpha^{3/2} \mathrm{e}^{-B\alpha}, \qquad B = \frac{X_c^2}{2\epsilon},$$

we set $\lambda(\alpha) \approx 1/t$:

$$C_1 \alpha^{3/2} \mathrm{e}^{-B\alpha} \approx \frac{1}{t}.$$

Taking the logarithm,

$$\ln C_1 + \frac{3}{2}\ln\alpha - B\alpha = -\ln t.$$

In the rare-event limit, $B\alpha \gg 1$, the exponential term dominates, giving

$$\alpha(t) \approx \frac{1}{B}\ln(C_1 t).$$

Therefore

$$\frac{\mathrm{d}\alpha}{\mathrm{d}t} = \frac{1}{Bt},$$

and

$$m(t) = P(\alpha)\left|\frac{\mathrm{d}\alpha}{\mathrm{d}t}\right| = \frac{1}{\alpha_{\max} Bt} = \frac{2\epsilon}{X_c^2\alpha_{\max}}\frac{1}{t}. \tag{S17}$$

##### S3.3 Uniform distribution of threshold distance $X_c$

Now assume a uniform distribution of threshold distances:

$$P(X_c) = \frac{1}{X_{c,\max}}, \qquad X_c \in [0, X_{c,\max}].$$

The escape rate can be written as

$$\lambda(X_c) = AX_c\exp(-BX_c^2), \qquad B = \frac{\alpha}{2\epsilon}, \qquad A = \sqrt{\frac{8\alpha^3}{\pi\epsilon}}.$$

Setting $\lambda(X_c) \approx 1/t$ gives

$$AX_c\mathrm{e}^{-BX_c^2} \approx \frac{1}{t}.$$

Taking the logarithm,

$$\ln A + \ln X_c - BX_c^2 = -\ln t.$$

In the rare-event regime, the quadratic barrier term dominates:

$$X_c(t) \approx \sqrt{\frac{1}{B}\ln(At)}.$$

Unlike the linear-potential case, where the frontier moves as $\ln t$, the harmonic well produces a slower frontier motion proportional to $\sqrt{\ln t}$. Differentiating,

$$\frac{\mathrm{d}X_c}{\mathrm{d}t} = \frac{1}{2\sqrt{B}}\frac{1}{t\sqrt{\ln(At)}}.$$

The mortality rate is therefore

$$\begin{aligned} m(t) &\approx \frac{1}{X_{c,\max}} \frac{1}{2\sqrt{B}} \frac{1}{t\sqrt{\ln(At)}} \\ &= \frac{1}{X_{c,\max}} \sqrt{\frac{\epsilon}{2\alpha}} \frac{1}{t\sqrt{\ln\left(\sqrt{\frac{8\alpha^3}{\pi\epsilon}}\, t\right)}}. \end{aligned} \tag{S18}$$

For $t \gg \sqrt{\pi\epsilon/(8\alpha^3)}$, the leading behavior is close to $1/t$, with a slowly varying logarithmic correction.

### S4 Summary of mortality scalings

Table S1 summarizes how the macroscopic mortality rate depends on the escape kinetics and on the distribution of barrier distances. Here $h(t)$ denotes the cohort hazard or mortality rate.

Table S1: Mortality scaling for different escape kinetics and barrier distributions.

| **Barrier distribution** $P(X_c)$ | **Linear potential,** $\lambda \sim \mathrm{e}^{-X_c}$ | **Quadratic potential,** $\lambda \sim \mathrm{e}^{-X_c^2}$ | **Inverse potential,** $\lambda \sim \mathrm{e}^{-1/X_c}$ |
|---|---|---|---|
| Uniform, $P \sim 1$ | $h(t) \sim 1/t$ | $h(t) \sim 1/[t\sqrt{\ln t}]$ | $h(t) \sim 1/[t(\ln t)^2]$ |
| Exponential decay, $P \sim \mathrm{e}^{-\gamma X_c}$ | $h(t) \sim t^{-(1+\gamma)}$ | $h(t) \sim \mathrm{e}^{-\gamma\sqrt{\ln t}}/[t\sqrt{\ln t}]$ | $h(t) \sim 1/[t(\ln t)^2]$ |
| Exponential growth, $P \sim \mathrm{e}^{+\gamma X_c}$ | $h(t) \sim t^{-(1-\gamma)}$ | $h(t) \sim \mathrm{e}^{+\gamma\sqrt{\ln t}}/[t\sqrt{\ln t}]$ | $h(t) \sim 1/[t(\ln t)^2]$ |
| Power-law decay, $P \sim X_c^{-k}$ | $h(t) \sim 1/[t(\ln t)^k]$ | $h(t) \sim 1/[t(\ln t)^{(k+1)/2}]$ | $h(t) \sim (\ln t)^{k-2}/t$ |
| Power-law growth, $P \sim X_c^{k}$ | $h(t) \sim (\ln t)^k/t$ | $h(t) \sim (\ln t)^{(k-1)/2}/t$ | $h(t) \sim 1/[t(\ln t)^{k+2}]$ |
